# The partitioning of symbionts effects on host resource acquisition and developmental plasticity

**DOI:** 10.1101/2020.04.27.064667

**Authors:** Robin Guilhot, Anne Xuéreb, Simon Fellous

**Affiliations:** CBGP, INRAE, CIRAD, IRD, Montpellier SupAgro, Univ Montpellier, Montpellier, France

## Abstract

Many symbionts provide nutrients to their host and/or affect its phenotypic plasticity. Such symbiont effects on host resource acquisition and allocation are often simultaneous and difficult to disentangle. Here we partitioned symbiont effects on host resource acquisition and allocation using a new framework based on the analysis of a well-established trade-off between host fitness components. This framework was used to analyze the effect of symbiotic yeast on the larval development of *Drosophila* larvae in field-realistic conditions. The screening of eighteen yeast fresh isolates showed they had similar effects on the resource acquisition in *Drosophila melanogaster, D. simulans* and *D. suzukii* but species-specific effects on resource allocation between either larval development speed or adult size. These differences shed light on the ecology of *Drosophila* flies and illustrate why distinguishing between these qualitatively different effects of microorganisms on hosts is essential to understand and predict symbiosis evolution.

## Introduction

Most macroorganisms are associated with symbionts that can provide nutrients (*i.e.*, resource acquisition thereafter) and/or affect their physiology and therefore their life-history traits (resource allocation hereafter) (Krajmalnik-Brown *et al.* 2012). Phenotype, and therefore fitness, usually depends on the amount of resource acquired by an organism (resource acquisition) and the proportion allocated to several dependent traits (resource allocation) (Van Noordwijk and De Jong 1986; De Jong and Van Noordwijk 1992). Individuals hence allocate limited amounts of energy to some traits at the detriment of others (Stearns 1989). Many examples of such trade-offs have been thoroughly documented in various organisms ranging from plants to microorganisms (Lind *et al.* 2013; Nørgaard *et al.* 2020). For example, lifespan and reproductive rate are constrained by a trade-off (Chippindale *et al.* 2004; Flatt 2011; Travers, Garcia-Gonzalez and Simmons 2015) because endogenous resources allocated to reproductive functions are not available for somatic and repair functions (Flatt, 2011; Schwenke, Lazzaro and Wolfner, 2016). Another allocation trade-off between speed of larval development and adult size has been largely described in arthropods: longer development time is costly but enables larger adult body size (Stearns and Koella 1986; Nunney 1996; DeLong and Hanley 2013; Teder, Vellau and Tammaru 2014). Despite the central role of trade-offs in life-history theory, positive correlations between traits that participate to fitness (*i.e.*, fitness components) are often observed in empirical datasets (Van Noordwijk and De Jong 1986; Metcalf 2016). These patterns occur when variation in resource acquisition is superior to variation in resource allocation. Indeed, organisms that acquire most resources can simultaneously maximize trait value in several fitness components, a phenomenon often referred to as ‘big car big house paradox’ (*e.g.*, Nunney 1996). When genes, or symbiont, simultaneously affect resource acquisition and allocation, partitioning between each type of effects may prove challenging.

Numerous microbial symbionts influence host resource allocation. For example, microbial symbionts affect hosts reproductive strategy along the lifespan - fecundity allocation trade-off in invertebrates (Emelianoff *et al.* 2008; Gould *et al*. 2018; Walters *et al.* 2019) and plants (Gundel *et al.* 2013; Yule, Miller and Rudgers 2013; Chung, Miller and Rudgers 2015). For instance, we recently reported bacterial symbionts of *Drosophila melanogaster* that also affect host developmental plasticity along the trade-off between speed of larval development and adult size (Guilhot *et al.* 2019). Other microbial symbionts mainly vary in their effects on host resource acquisition. In that case, as for ‘the big car big house paradox’, the screening of several symbiont strains may reveal the simultaneous maximization of several fitness components. This was also reported with *D. melanogaster* symbionts (*e.g.*, Anagnostou, Dorsch and Rohlfs 2010; Bing *et al.* 2018; Pais *et al.* 2018). Most studies have so far concluded that symbionts affect either host resource acquisition or allocation. However, there is no reason symbionts would not simultaneously affect both type of variation. Here we introduce of new framework that enables to partition symbiont effects on host resource acquisition and allocation, and we illustrate its relevance with an experiment on *Drosophila*-yeast symbiosis in field-realistic conditions.

*Drosophila* flies are associated with nutritional extracellular symbionts that provide nutrient to the host and affect its physiology (Vega and Dowd 2005; Shin *et al.* 2011; Storelli *et al.* 2011; Wong, Dobson and Douglas 2014; Sannino *et al.* 2018). Larval speed of development and adult body size are common measures of symbiont effects on *Drosophila* phenotypes (Anagnostou, Dorsch and Rohlfs 2010; Bellutti *et al.* 2018; Bing *et al.* 2018; Lewis and Hamby 2019; Murgier *et al.* 2019). These two traits, that are constrained by a trade-off (Teder, Vellau and Tammaru 2014) are both correlated positively with fitness in numerous insects (Preziosi *et al.* 1996; Nylin and Gotthard 1998). We revealed the effects of yeast on resource allocation in *Drosophila* by investigating symbiont effects along the trade-off between speed of larval development and adult size. Symbiont effects on resource acquisition were revealed by yeast-induced variations of host phenotype orthogonal to the trade-off function (Figure 2).

## Material and methods

### Biological material

Eighteen yeast strains were used in this study (Table 1). Among them, sixteen strains were isolated from wild adults *Drosophila melanogaster* and *D. suzukii* and from grape flesh and skin samples collected near Montpellier, southern France in the late 2017. They belong to ten yeast and yeast-like taxa previously reported present in wild Drosophilids and their environment. Species included *Hanseniaspora uvarum*, a yeast that frequently associates with *D. suzukii* and *D. melanogaster* (Anagnostou, LeGrand and Rohlfs 2010; Barata, Malfeito-Ferreira and Loureiro 2012; Hamby *et al.* 2012; Hoang, Kopp and Chandler 2015; Bellutti *et al.* 2018), of which we added two other strains of reference. Molecular identification of each strain was carried out by sequencing of the ITS1 region (ITS1-F and ITS2 primers) of nuclear ribosomal RNA gene (White *et al.* 1990; Gardes and Bruns 1993).

**Table 1.** Yeast strains used in this study.

| Strain | Origin | Accession number | Sequence length | Closest strain | % Identity |
| --- | --- | --- | --- | --- | --- |
| Saturn. div. 1<br>(*Saturnispora diversa*) | *D. suzukii*, France | MN684811 | 150 | *Saturnispora diversa*<br>(KY105316) | 100.00 % |
| Cand. stell. 1<br>(*Candida stellimalicola*) | *D. melanogaster*,<br>France | MN684812 | 179 | *Candida stellimalicola*<br>(KP131811) | 96.65 % |
| Cand. stell. 2<br>(*Candida stellimalicola*) | *D. melanogaster*,<br>France | MN684813 | 179 | *Candida stellimalicola*<br>(KP131811) | 96.65 % |
| Star. bacill. 1<br>(*Starmerella bacillaris*) | *D. suzukii*, France | MN684814 | 198 | *Starmerella bacillaris*<br>(KY102529) | 100.00 % |
| Trigo. vin. 1<br>(*Trigonopsis vinaria*) | *D. melanogaster*,<br>France | MN684815 | 203 | *Trigonopsis vinaria*<br>(KY105765) | 100.00 % |
| Trigo. vin. 2<br>(*Trigonopsis vinaria*) | *D. suzukii*, France | MN684816 | 203 | *Trigonopsis vinaria*<br>(KY105765) | 100.00 % |
| Vishniaco. 1<br>(*Vishniacozyma sp.*) | *D. suzukii*, France | MN684817 | 211 | *Vishniacozyma carnescens*<br>(MG250423) | 100.00 % |
| Vishniaco. 2<br>(*Vishniacozyma sp.*) | Grape berry,<br>France | MN684818 | 211 | *Vishniacozyma carnescens* (KY105819) | 100.00 % |
| Rhodoto. 1<br>(*Rhodotorula sp.*) | Grape berry,<br>France | MN684819 | 228 | *Rhodotorula sp.*<br>(MH380198) | 100.00 % |
| Rhodoto. 2<br>(*Rhodotorula sp.*) | Grape berry,<br>France | MN684820 | 228 | *Rhodotorula sp.*<br>(MH380198) | 100.00 % |
| Aureobas. 1<br>(*Aureobasidium sp.*) | *D. suzukii*, France | MN684821 | 255 | *Aureobasidium sp.*<br>(MH399294) | 100.00 % |
| Aureobas. 2<br>(*Aureobasidium sp.*) | Grape berry,<br>France | MN684822 | 255 | *Aureobasidium sp.*<br>(MH399294) | 100.00 % |
| Sacchar. crata. 1<br>(*Saccharomycopsis crataegensis*) | *D. suzukii*, France | MN684823 | 266 | *Saccharomycopsis crataegensis*<br>(KY105255) | 100.00 % |
| Hans. mey. 1<br>(*Hanseniaspora meyeri*) | *D. suzukii*, France | MN684825 | 363 | *Hanseniaspora meyeri*<br>(KY103535) | 100.00 % |
| Hans. uv. 1<br>(*Hanseniaspora uvarum*) | *D. melanogaster*,<br>France | MN684824 | 364 | *Hanseniaspora uvarum*<br>(KY103571) | 100.00 % |
| Hans. uv. 2<br>(*Hanseniaspora uvarum*) | Grape berry,<br>France | MN684826 | 333 | *Hanseniaspora uvarum*<br>(KY103573) | 99.70 % |
| Hans. uv. 3<br>(*Hanseniaspora uvarum*) | *D. suzukii* | not<br>referenced |  |  |  |
| Hans. uv. 4<br>(*Hanseniaspora uvarum*) | *Drosophila sp.* | KF958056.1 |  |  |  |

#### The *Drosophila* groups used in this study were composed of

A *D. melanogaster* – *D. simulans* group, composed of two *Drosophila melanogaster* populations: an OregonR strain (in laboratories since 1927) and a strain we named Grenade as it was founded from a few dozen of adults that emerged from a pomegranate fruit (*i.e.*, Grenade in French) collected 6 months before the experiment near Montpellier, Southern France; and one *D. simulans* population (Watsonville, founded four years before the experiment, from wild individuals collected in California, USA).

A *D. suzukii* group, composed of two *D. suzukii* populations: Watsonville, founded four years before the experiment, from wild individuals collected in California, USA; and GaMu, founded 5 years before the experiment, from wild individuals collected in Gaujac, southern France).

These fly populations have been maintained in the laboratory at 21°C, 70% humidity and a 14 h photoperiod on a carrot-based laboratory medium (37.5 g.L^-1^ sugar, 37.5 g.L^-1^ dried carrot powder (Colin Ingredients SAS), 22.5 g.L^-1^ inactive dry yeast, 15 g.L^-1^ corn meal, 11.25 g.L^-1^ agar, 5 mL.L^-1^ propionic acid, 3.3 g.L^-1^ nipagin, 2.5 mL.L^-1^ ethanol).

### Experimental design

We investigated the effects of each yeast strain on the traits of each fly population on real fruit. Grape berries were sterilized in surface using successive baths of soap, bleach, ethanol and sterile water according to the protocol of Behar *et al.* (2008). One experimental unit was constituted of a halved sterilized fruit fixed at the center of a small petri dish with jellified sterile water. Ten eggs were deposited on the fruit surface. These eggs were obtained from grape juice agar plates exposed to fly females (300 mL.L^-1^ grape juice, 6 g.L^-1^ agar supplemented with 1 mg.L^-1^ cycloheximide to suppress extracellular yeast - but not bacteria - from the egg chorions). After the egg deposit, 10^2^ cells of a yeast strain (from 50 µL of yeast stock conserved in PBS + 20% glycerol at -80°C) were inoculated on the fruit surface (Figure 1A). We created six to seven replicates of each yeast strain x fly population combination and six to ten replicates of controls (larvae of each population without yeast) over seven experimental blocks. Petri dishes were placed in a climatic chamber at 24°C. Once larvae transform into pupae (Figure 1B), petri dishes were controlled daily to collect adults in 96° ethanol (Figure 1C). Adult body size was estimated using thorax size, which was measured from the most anterior margin of the thorax to the posterior tip of the scutellum using a binocular microscope (Figure 1D) (Loeschcke, Bundgaard and Barker, 2000).

**Figure 1.**
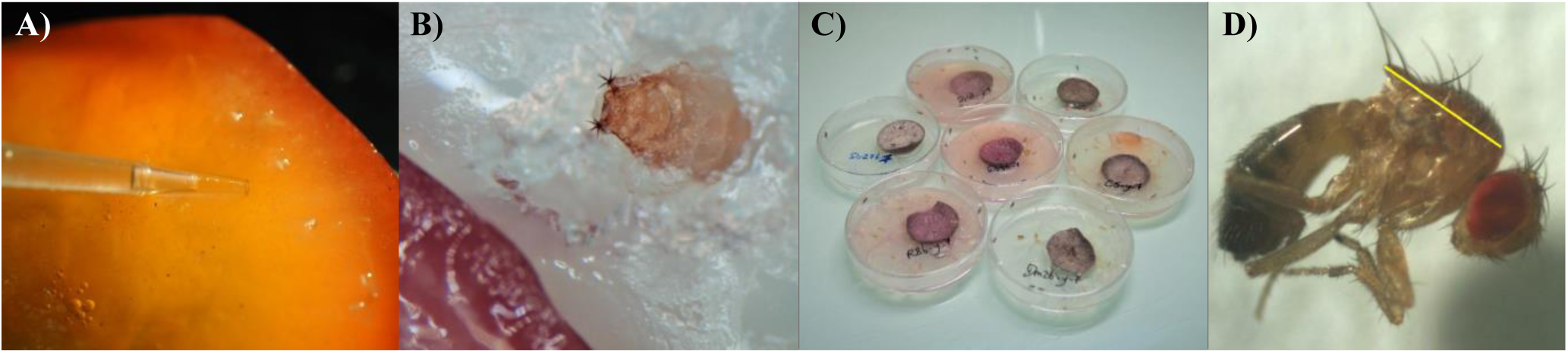
Experimental protocol. (A) Inoculation of yeast cells around *Drosophila* eggs. (B) Pupation on the fruit or the surrounding environment. (C) Experimental units with freshly emerged adults. (D) Estimation of thorax size (photography from Chechi *et al.* (2017)).

**Figure 2.**
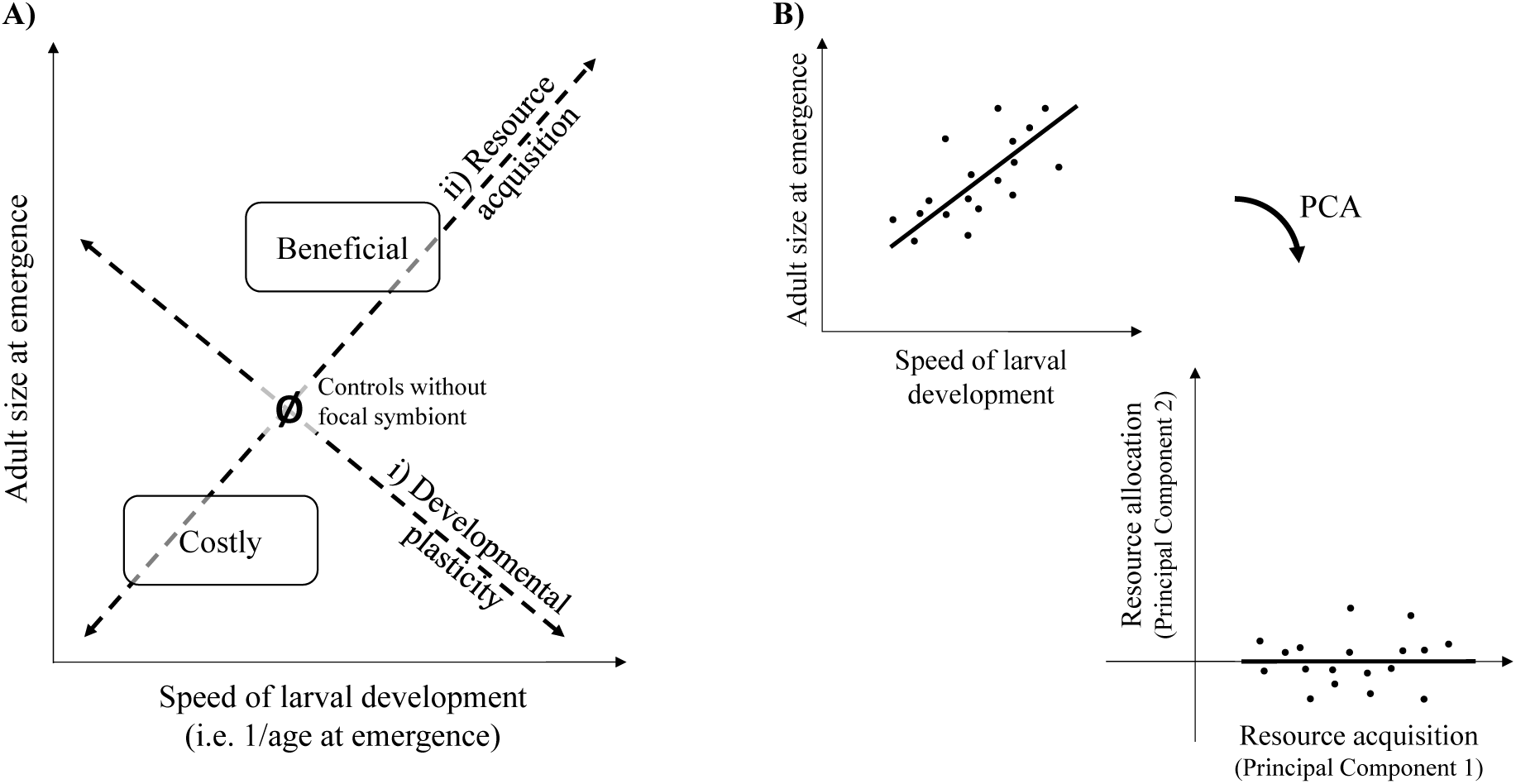
Analytical framework. (A) Phenotypic space defined by host larval speed of development and size at emergence. Host phenotypes induced by symbionts vary along i) the allocation trade-off (*i.e.*, developmental plasticity) axis and ii) the resource acquisition axis (defined as orthogonal to the allocation trade-off axis). Symbionts that induced faster development time and / or bigger individuals than control individuals (symbolized by Ø) are considered as beneficial while symbionts that induced slower development time and smaller individuals than control individuals are considered as costly (Stearns and Koella 1986; Jaramillo, Mehlferber and Moore 2015). (B) Position of datapoints after rotation of the phenotypic space.

To estimate yeast density in fruit flesh, all fruit substrates from two different blocks were sampled using five sterile pipette tips six to seven days after the petri creation. Samples were homogenized in sterile PBS solution, serially-diluted and plated on YPD (Yeast Extract-Peptone-Dextrose) agar supplemented with 10 mg.L^-1^ chloramphenicol incubated at 24°C for two – four days.

Growth of exogenous microorganisms to our study was observed in 17.08% of the replicates. We did not exclude these replicates from our analyses as these microorganisms, that could possibly be endophytes (Schulz and Boyle, 2005), did not expressed randomly among yeast treatments and did affect fly traits (Supplementary Material 1).

## Statistical analyses

### Overall reasoning

We followed a four steps path to analyze yeast effect on fly development. Our strategy was to first (1) get an overview of yeast effects on each of the single phenotypic traits we collected. We (2) studied the relationship between speed of larval development and adult size. This analysis revealed that yeast generally improved both traits, indicative of substantial resource provisioning by the symbiont. In order to disentangle yeast effects on resource acquisition and allocation (*i.e.*, developmental plasticity along the speed of development – adult size trade-off), we (3) constructed two new composite variables describing resource acquisition and allocation. Eventually, we (4) investigated how yeast strain features (*i.e.*, cell number and origin when sampled) influenced host life-history traits.

### Analysis of yeast effects on individual traits

To investigate the effects of yeast treatment on each individual trait, proportion of eggs that survived until adult emergence in each fruit, adult age at emergence (*i.e.*, 1/age at emergence) and thorax size were measured, and linear mixed models (REML method) were used for the analysis. Initial models included ‘*yeast treatment*’ (*i.e.*, all tested strains and the controls), ‘*fly group*’ (*i.e., D. suzukii* or *D. melanogaster* and *D. simulans* together), ‘*fly population*’ (nested in ‘*fly group*’) and their interactions all defined as fixed factors. Models also contained ‘*experimental block*’ as random term, and ‘*experimental unit*’ in the case of adult size analysis (*i.e.*, to control for the fact that we measured several adults per infested fruit). Because it is well established that yeast presence improves *D. melanogaster* larval survival we contrasted treatments with and without yeast in a similar model. Here, and everywhere else in the study, we followed backward stepwise model selection procedures so as to eliminate non-significant terms from the initially complex models.

In all cases, and later on in this manuscript, we ensured homoscedasticity and residuals normality complied with model assumptions. All analyzes were performed with JMP (SAS, 14.1). The dataset will be available in the open data repository Zenodo (DOI: to be determined).

### Simultaneous effects of yeast on speed of larval development and adult size

In a first attempt to describe how symbionts affect larval speed of development and adult thorax size, we plotted the means of each yeast treatment in this bi-dimensional phenotypic space. It was necessary to use the mean phenotypic value per treatment to avoid variations among replicates of the same treatment. These variations would have reflected micro-environmental variations (*e.g.*, fruit quality) rather than overall effects of the yeast strains tested. In these analyses, we used a single data point per combination of yeast treatment, fly population and fly sex. Based on prior work (Guilhot *et al.* 2019), we expected that a negative correlation between these two traits would indicate that yeasts affected mostly host resource allocation (Figure 2), while a positive would reflect substantial variation in host resource acquisition among the yeast strains. We performed two regression methods: the asymmetrical ordinary least squares regression (OLS), which enables to test differences among fly populations, and the symmetrical orthogonal regression (reported in Supp Mat).

### Constructing new composite variables describing yeast effects on resource acquisition and allocation

The analysis of yeast effect based on mean traits per treatment suggested that symbionts simultaneously impact resource acquisition and allocation. In order to disentangle yeast effects on each aspect of host physiology, we constructed a new composite variable describing each of them independently. We followed a method inspired from the original framework of Van Noorwijk and De Jong (1986) and the more recent literature on intra-locus sexual conflict (*e.g.*, Berger *et al.* 2014). Namely, we assumed that a positive covariation between fitness components (here speed of larval development and adult size) reflects variations in resource acquisition; variation along a trade-off between fitness component reflected resource allocation variations. This amounted to rotating the phenotypic space defined by speed of development and adult thorax size (Figure 2B). We explored several approaches, which all converged, to define these axes in our dataset, using either all datapoints or means per treatments. Eventually, we used a principal component analysis (PCA) to rotate the phenotypic space: resource acquisition was indicated by the first axis and resource allocation by the second (Fig SM4). Note that this solution was possible because our dataset was mostly structured by resource acquisition variation, which saved us from defining the slopes of composite axes *a priori*. One PCA per fly group and per sex was performed with all datapoints. Rotations per sex and fly species were mandatory as each of them placed in different areas of phenotypic space: in a joint PCA major variation axes would have been driven by sex- and species-induced variations on resource allocation rather than acquisition. Note that rotations per fly population were impossible due to a lack of positive relationship between speed of development and adult thorax size for the *D. suzukii* W population (see Results section).

Eventually, we carried out the same type of analysis on each of the new composite axes as for single trait analyses. In brief, we used linear mixed models to investigate yeast and fly influence resource acquisition and allocation.

### How yeast features do determine their effect on host phenotype?

We investigated how yeast features, namely microbial cell numbers in fruit flesh and the host species the strains were associated to when sampled (*i.e.*, yeast origin hereafter), impact host life-history traits. We hypothesized that yeast strains that multiplied the most would be associated with a greatest resource acquisition, as would do yeasts sampled in the same host species than the one they were tested in. We used the same type linear mixed models as above to analyze survival, development time, size at emergence, resource acquisition and resource allocation. However, we excluded yeast-free controls from the analysis of cell numbers and controls as well as the two laboratory strains of *H. uvarum* from the yeast origin analysis.

Effects of yeast cell numbers and origin were therefore tested in separate models. In the first, we added log-transformed cell numbers as covariate to models. In the second, we added a yeast origin term within which yeast treatment was nested. Interactions with these terms were also tested. Eventually, we studied with a linear mixed model the factors determining variation in yeast cell numbers.

## Results

### Yeast symbionts affected differently individual traits of two host groups

Developmental survival, *i.e.*, the proportion of eggs surviving until the adult stage, responded differently to yeast inoculation in the two fly groups (F-test of ‘*presence/absence of yeast*’ *\** ‘*fly group*’: F_1,626_ = 8.27, p = 0.0042) (Figure 3A). Yeast inoculation enhanced the survival of *D. melanogaster* and *D. simulans* larvae (contrast ‘*yeast presence*’ vs ‘*control’*: F_1,628_ = 25.21, p < 0.0001), but had no effect on *D. suzukii* (contrast ‘*yeast presence*’ vs ‘*control*’: F_1,625_ = 0.03, p = 0.8547). Furthermore, developmental survival was influenced by yeast treatment but not by interactions between this factor and the fly identity; in other words symbionts had overall a similar effect on all hosts (Table 2, Figure 3B).

**Table 2.** Analysis of individual phenotypic traits. Linear mixed models (REML), final models excluding non-significant terms.

|  | Developmental survival | Age at emergence | Thorax size |
| --- | --- | --- | --- |
| Yeast treatment | $F_{18,591} = 3.98; p < 0.0001$ | $F_{18,559} = 23.38; p < 0.0001$ | $F_{18,540} = 6.45; p < 0.0001$ |
| Fly population | $F_{3,592} = 10.15; p < 0.0001$ | $F_{3,544} = 21.92; p < 0.0001$ | $F_{3,523} = 9.44; p < 0.0001$ |
| Fly group | $F_{1,594} = 34.78; p < 0.0001$ | $F_{1,566} = 253.84; p < 0.0001$ | $F_{1,517} = 2404; p < 0.0001$ |
| Yeast treatment * fly group | $F_{18,591} = 1.42; p = 0.1157$ | $F_{18,559} = 2.20; p = 0.0031$ | $F_{18,540} = 1.70; p = 0.0363$ |
| Sex | | $F_{1,3344} = 2.11; p = 0.1468$ | $F_{1,2608} = 3195; p < 0.0001$ |

**Figure 3.**
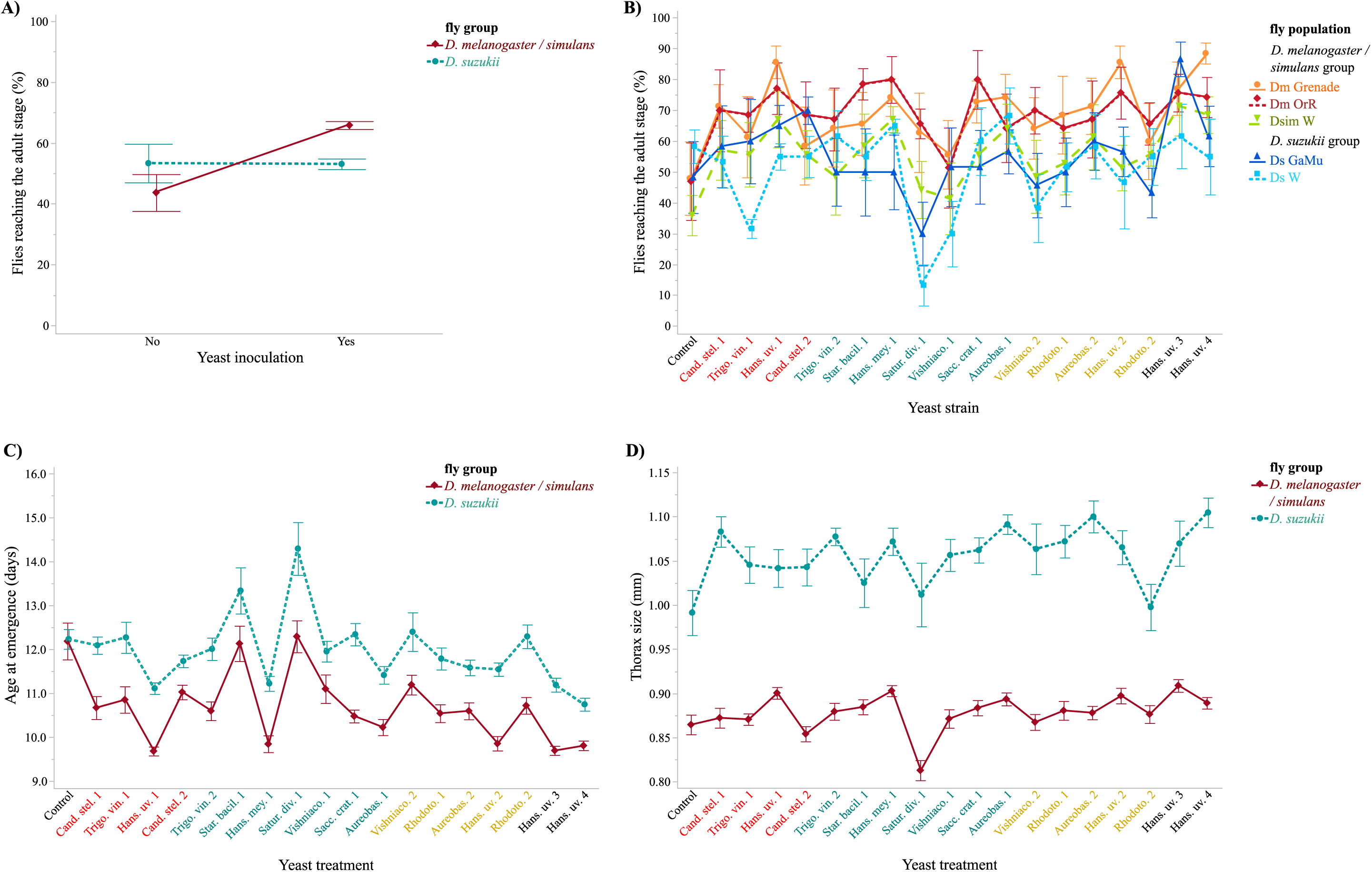
Effects of yeast symbionts on fly individual traits. (A) Developmental survival of each fly group in absence or presence of yeast. (B) Developmental survival of each fly population per yeast treatment. (C) Age at emergence of each fly group per yeast treatment. (D) Thorax size of each fly group per yeast treatment. Symbols indicate means and error bars indicate standard errors around the mean. X axis color code indicates the origin of the strains (red: *D. melanogaster*, blue: *D. suzukii*, green: fruit). Additional, more detailed figures (grouping per fly population) are presented in Supplementary Material 2.

Age at emergence and adult thorax size were both significantly influenced by an interaction between the yeast treatment and the fly group. Yeast strains did not have different effects males and females (Table 2, Figures 3C and 3D).

### Simultaneous and beneficial effects of yeast on larval speed of development and adult size

In *D. melanogaster* and *D. simulans* flies, relationships between effects of yeasts on larval speed of development and adult thorax size at emergence were always positive and significant (Figure 4, Table 3). In *D. suzukii* group, the relationships were either marginally significant and positive (Ds GaMu males and females) or non-significant (Ds W males and females) (Figure 4, Table 3). In vast majority of cases, yeast strains led to insect hosts that were both bigger and faster to develop than yeast-free control (Figure 3, controls represented with solid symbols), suggestive of substantial nutrient provisioning of the host by the symbionts. We interpreted the non-significant regression in *D. suzukii* as indicative that in addition to effects on resource acquisition, yeasts also impacted the developmental plasticity of these insects along the trade-off between speed of larval development and adult size. Indeed, this variation would have spread datapoints away from the regression axis reducing consequently the fit of the model.

**Table 3.**
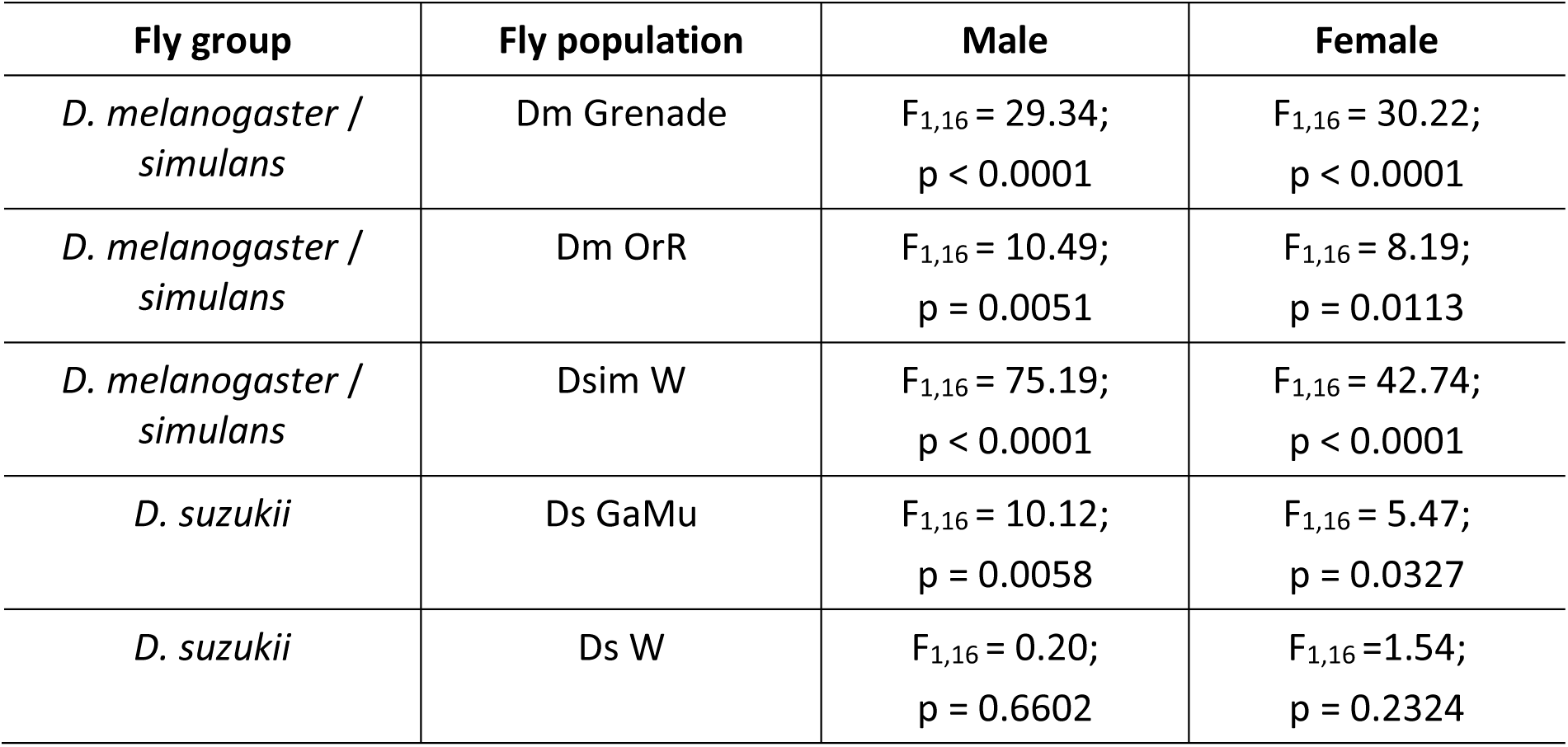
Linear regressions of adult thorax size onto larval speed of development.

| Fly group | Fly population | Male | Female |
| --- | --- | --- | --- |
| *D. melanogaster / simulans* | Dm Grenade | $F_{1,16} = 29.34;$<br>$p < 0.0001$ | $F_{1,16} = 30.22;$<br>$p < 0.0001$ |
| *D. melanogaster / simulans* | Dm OrR | $F_{1,16} = 10.49;$<br>$p = 0.0051$ | $F_{1,16} = 8.19;$<br>$p = 0.0113$ |
| *D. melanogaster / simulans* | Dsim W | $F_{1,16} = 75.19;$<br>$p < 0.0001$ | $F_{1,16} = 42.74;$<br>$p < 0.0001$ |
| *D. suzukii* | Ds GaMu | $F_{1,16} = 10.12;$<br>$p = 0.0058$ | $F_{1,16} = 5.47;$<br>$p = 0.0327$ |
| *D. suzukii* | Ds W | $F_{1,16} = 0.20;$<br>$p = 0.6602$ | $F_{1,16} = 1.54;$<br>$p = 0.2324$ |

**Figure 4.**
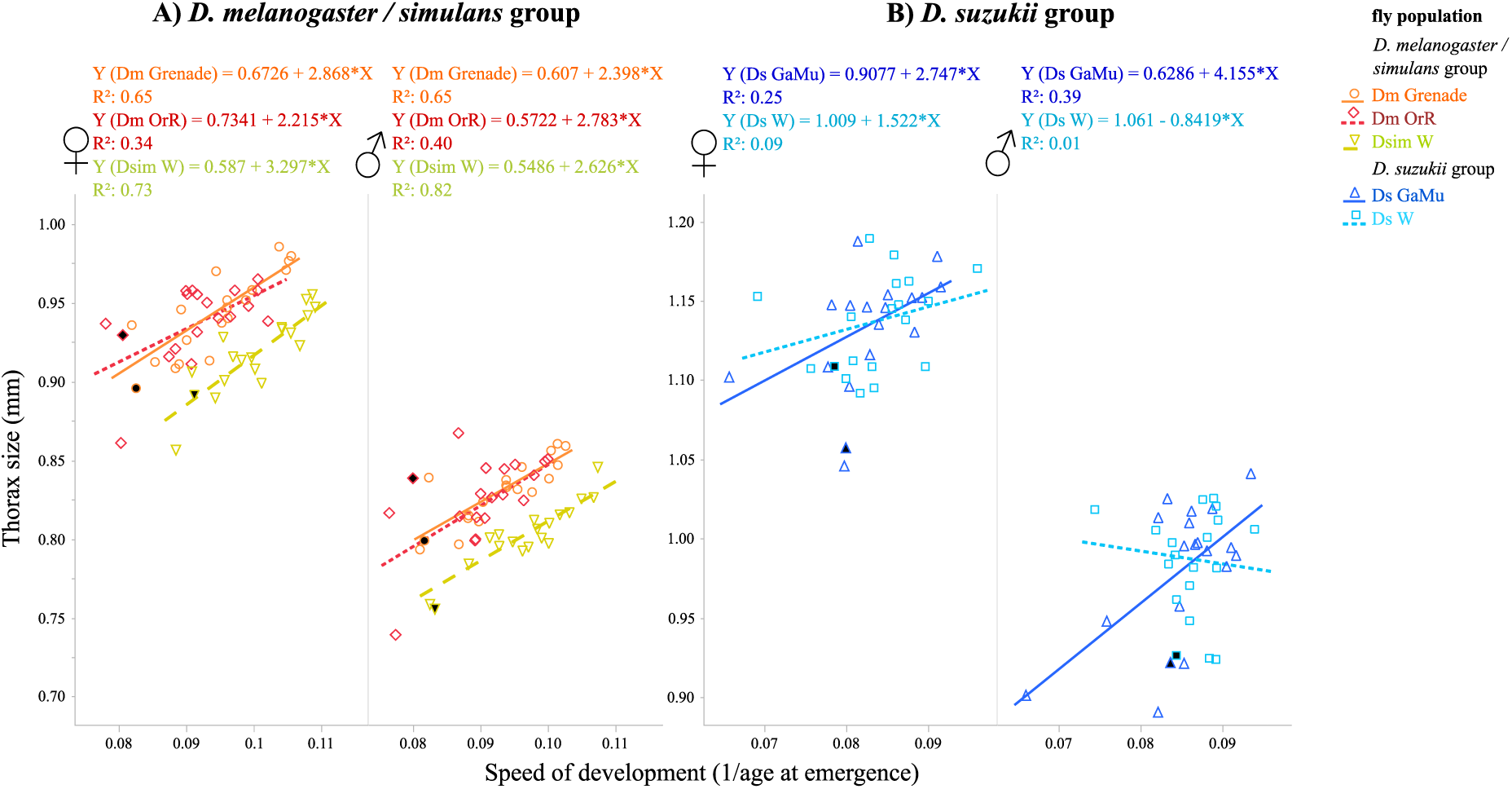
Relationships between yeast effects on larval speed of development and adult thorax size. Each point in the phenotypic space represents means of the two life-history traits for each yeast strain in each fly population and sex. Control phenotypes (*i.e.*, without yeast symbionts) are indicated by black-filled symbols. Ordinary least-squares are presented here, symmetrical regression are presented in SM 3.

### Partition of yeast influence into resource acquisition and allocation effects

After phenotype space rotation, and therefore the computation of new composite variables describing symbiont effects on host resource acquisition and allocation, we deployed the same type of analyses as for the original traits. Resource acquisition was largely determined by yeast treatment main effect: yeast strains varied in their boosting effects on fly hosts irrespective of their species, population or sex (Figure 5A, Table 4). However, host resource allocation was determined by a largely significant interaction between the yeast treatment and fly group. In other words, the different yeast strains had different effects on the developmental plasticity of *D. melanogaster* / *simulans* compared to that of *D. suzukii* (Figure 5B, Table 4). Eventually, we tested whether effects of yeasts on host resource allocation were more variable in *D. suzukii* than in *D. melanogaster* and *simulans*, which was confirmed by a non-parametric Levene test (F_1,34_= 6.8, p= 0.013, tested on means per yeast strain and fly group).

**Table 4.** Analysis of resource acquisition and allocation. Linear mixed models (REML), final models excluding non-significant terms.

|  | Resource acquisition (PC1) | Resource allocation (PC2) |
| --- | --- | --- |
| Yeast treatment | $F_{18,551} = 20.85$; $p < 0.0001$ | $F_{18,557} = 6.37$; $p < 0.0001$ |
| Fly population [Fly group] | $F_{3,540} = 2.85$; $p = 0.0367$ | $F_{3,544} = 53.96$; $p < 0.0001$ |
| Fly group | $F_{1,559} = 1.02$; $p = 0.3122$ | $F_{1,567} = 0.40$; $p = 0.5276$ |
| Yeast treatment * fly group | $F_{18,552} = 1.49$; $p = 0.0880$ | $F_{18,557} = 2.88$; $p < 0.0001$ |

**Figure 5.**
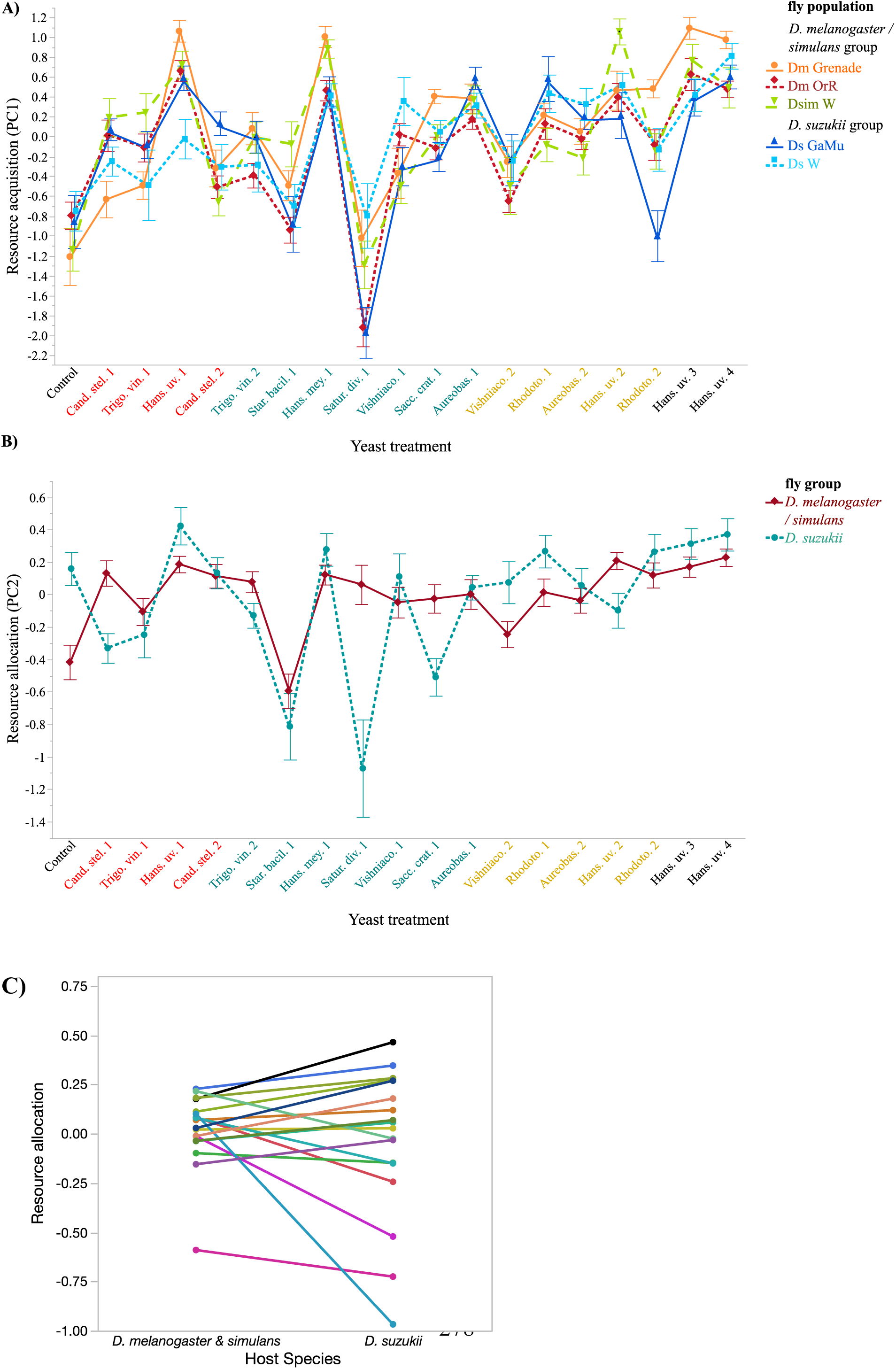
Fly resource acquisition and allocation in response to yeast treatment. (A) Fly resource acquisition in response to yeast treatment. (B) Fly resource allocation in response to yeast treatment (an additional figure is presented in Supplementary Material 4). C) Effects of yeasts strains on host resource allocation were more variable in *D. suzukii* than in *D. melanogaster* and *simulans* (Levene test, F_1,34_= 6.8, p= 0.013). Symbols indicate means and error bars indicate standard errors around the mean. X axis color code indicates the origin of the strains (red: *D. melanogaster*, blue: *D. suzukii*, green: fruit).

### Determinants of yeast effects on host phenotypes

The different yeast strains harbored very different in-fruit cell numbers (F-test of ‘*yeast treatment*’: F_17,147_ = 4.39; p < 0.0001, Figure S6.1). However, yeast cell numbers had no significant effects on either host resource acquisition or allocation. Accordingly, cell numbers did not correlate with any of the original or composite traits we studied (Figure S6.2).

The host species from which yeast strains had been isolated (*i.e.*, yeast origin) had significant effects on both resource acquisition and allocation. A remarkable interactive effect between yeast origin and fly group on resource allocation was due to the response of *D. suzukii* flies associated to yeast isolated from the same species (contrast “*D. suzukii hosts with D. suzukii yeast*” vs “*all other combinations*”: F_1, 483_= 23.75; p<0.0001). Indeed, this combination produced insects that positioned on the faster but smaller side of the developmental plasticity trade-off (Figure 6).

**Figure 6.**
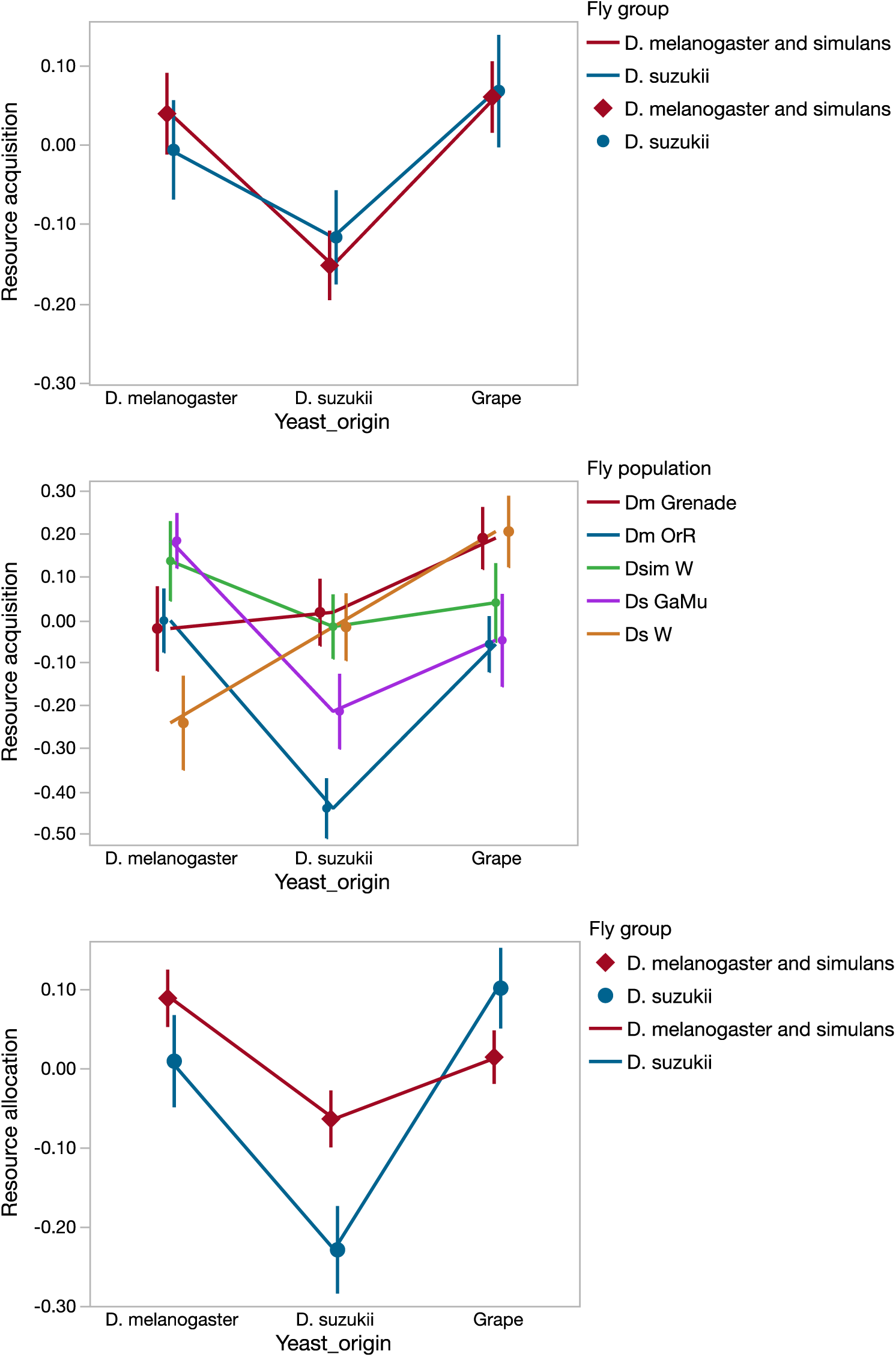
Effect of yeast origin on resource allocation and acquisition. Symbols indicate means and error-bars standard-errors.

## Discussion

We studied the influence of eighteen strains of symbiotic yeast on the larval development of *Drosophila* flies in field-realistic conditions. A new analytical framework based on the joint analysis of traits constrained by a physiological trade-off (speed of larval development and adult size) enabled to distinguish the effect of symbionts on host resource acquisition from those on resource allocation (*i.e.*, provisioning and developmental plasticity, respectively). If yeast had similar influence on resource acquisition in all fly species tested, their effects on resource allocation drastically different among host species. In particular, we found that yeast strain diversity led to greater differences in resource allocation in *D. suzukii* than in *D. melanogaster* and *D. simulans*. Besides, the host of origin of the yeast strains (*i.e., D. suzukii, D. melanogaster* or grape) affected resource allocation in a host species specific fashion, which was not the case for resource acquisition. None of these effects were visible when analyzing host traits individually.

### Partition symbiont effects to unveil patterns of evolutionary importance

The framework we developed here aimed at distinguishing symbiont effects on host resource acquisition and allocation. It was motivated by the different evolutionary consequences of symbiont effects on these two types of functions (see below). Our framework revealed that symbiotic yeast strains greatly varied in terms of resource provisioning, but these effects remained similar on all host populations and species tested. By contrast, yeast influence on host resource allocation (*i.e.*, its developmental plasticity along the trade-off between speed of larval development and adult size) differed when symbionts were associated to *D. suzukii* or *D. melanogaster* and *D. simulans* (Figures 5 and 6). Previous work reported that symbionts can affect host resource allocation, even in the *Drosophila* system (Gould *et al.* 2018; Guilhot *et al.* 2019; Walters *et al.* 2019). To our knowledge, however, none had partitioned the simultaneous effects of symbionts on resource acquisition and allocation. It may appear surprising as it has been known for several decades that each is under simultaneous genetic influence and that discriminating between acquisition and allocation of resource has been an important advancement in life-history theory (Van Noordwijk and De Jong 1986; De Jong and Van Noordwijk 1992). Nutritional symbionts are often described either to affect host nutrient provisioning or to influence host physiology through a modulation of nutrient allocation (Krajmalnik-Brown *et al.* 2012). These two mechanisms can have drastically different influences on the evolution of host – microorganism symbioses (Fisher *et al.* 2017; Brown and Akçay 2019; Moran, Ochman and Hammer 2019). For example, symbionts that provide valuable nutrients to the host would improve host fitness in most environmental contexts, which may lead to stable mutualisms and even evolved dependence (Fellous and Salvaudon 2009). By contrast, symbionts that mostly affect host developmental plasticity will induce host phenotypes that may or may not be adequate to current environmental conditions. Their costs and benefits will depend on the environment, which may lead to conditional mutualisms. As a consequence, symbionts that modulate host development and plasticity would enable alternative host life-history strategies and participate most to local adaptation (Metcalf *et al.* 2019). Besides, mathematical modelling focusing on the reproduction-survival trade-off shows that whether symbionts affect one or the other trait can affect the evolution of the symbiont mode of transmission and therefore the evolutionary fate of the partnership (*e.g.*, Brown and Akçay 2019). It was recently argued that different bacterial communities participate to *D. melanogaster* local adaptation in the field (Walters *et al.* 2019). Along these lines, we discuss below how the patterns unveiled by our framework make sense (in evolutionary terms) in the light of the ecology of the hosts.

### Ecological and physiological relevance of symbiont effects

One remarkable pattern our framework unveiled was the different variance of yeast strains effects on host resource allocation in the two fly groups (Figure 5C). In *D. suzukii*, yeast strain had a considerable influence on the developmental pathway followed by the host along the fast but small – slow but large trade-off. By comparison, the development of *D. melanogaster* and *simulans* flies was less sensitive (concerning the change in development) to the nature of the yeast strain they associated with. The phenomenon is reminiscent of environmental canalization where organisms produce similar phenotypes in different environments (*i.e.*, low phenotypic plasticity) (Flatt 2005). Responding excessively to environmental factors can indeed be detrimental, in particular when fitness landscapes are steep and fitness maximized in narrow ranges of trait values, or when organism experience unpredictable and highly variable developmental conditions. The latter is the case of *D. melanogaster*, which larvae usually develop in rotting fruits that contain a large diversity of microorganisms (*e.g.*, Stamps *et al.* 2012). *D. suzukii* females however prefer to oviposit in ripening fruit devoid of rot; the diversity of yeast strains their larvae are exposed to is therefore low (Hamby and Becher 2016). It is therefore possible that the lower developmental sensitivity of *D. melanogaster* and *D. simulans* to yeast strain identity is an adaptation to their saprophytic life-style. An alternative explanation may be that *D. suzukii* uses yeast symbionts more than *D. melanogaster* to manipulate its developmental trajectory. It is indeed well established *Drosophila* larvae and adults exert active choices resulting in their association with particular symbionts (*e.g.*, Wong *et al.* 2017).

Yeast strains sampled in different hosts had different effects on host resource allocation. These effects depended on the host species: yeast that came from *D. suzukii* accelerated *D. suzukii* larval development (at the cost of producing smaller adults). It is difficult to discuss this pattern without leaning towards spandrel explanation *à la* Gould and Lewontin (Gould and Lewontin 1979). One may however notice that a fast but small strategy (*i.e.*, an *r-* type strategy) maximizes fitness for population in exponential expansion, which is common of *D. suzukii* populations in orchards.

Yeast strains varied considerably in terms of host of resource provisioning (*i.e.*, their effect of host resource acquisition, Figure 5A), a frequent observation in *Drosophila* symbiosis (Anagnostou, Dorsch and Rohlfs 2010; Bing *et al.* 2018). However, we were surprised that yeast cell numbers did not correlate with any of the host traits measured (Figure S6.2). This result does not only contradict previous results obtained with bacteria in artificial conditions (Keebaugh *et al.* 2018), but also questions the nature of the resource provided by these symbionts to their hosts. Indeed, would yeast cells be mere calories, one would expect beneficial effects to correlate with in-fruit cell numbers. Yeast strains did vary in terms of cell multiplication; the most proficient ones were c.30 fold more numerous than the least proficient ones. Besides, cell dimensions of our yeast species are known to be all in the same range (Kurtzman, Fell and Boekhout 2011). We therefore conclude that the yeast strains studied mainly varied in the qualitative nature of the resources they provided. In fruit tissues, nutrients are typically enclosed in cells and numerous micronutrients are limiting (Vega and Dowd 2005). But different yeasts species produce different proteins, lipids (sterols, fatty acids) and micronutrients (amino acids, vitamins, mineral salts) due to their different metabolisms (Flores *et al.* 2000; Fanson and Taylor 2012). Yeast quality hence affects fly traits as demonstrated in *D. melanogaster* (*e.g.*, Grangeteau *et al.* 2018). The fact that yeast strains had similar effect on resource acquisition in all fly species was however unexpected. As discussed above, larvae of the two fly groups are thought to have different relationships with microbial symbionts: *D. melanogaster* and *D. simulans* larvae develop in rotten substrates rich in proteins while *D. suzukii* larvae develop on ripening fruits poorer in proteins (Capy and Gibert 2004; Lee *et al.* 2011; Lewis and Hamby 2019). As expected, the presence of yeasts did improve *D. melanogaster* and *D. simulans* larval survival until adult emergence and had no effect in *D. suzukii*, suggesting that the latter species is better adapted to poor media and confirming that most microorganisms enable the development of *D. suzukii* larvae in fruit (Bing *et al.* 2018).

While yeast apparently provisioned equally beneficial micronutrients to all species (Figure 5A), how each species exploited them differed. In presence of the symbiont, *D. suzukii* responded by increasing adult size (Figure 4B) whereas *D. melanogaster* and *D. simulans* accelerated larval development. Partitioning symbiont effects into resource acquisition and allocation revealed a number of phenomena impossible to quantify with the analysis of single phenotypic traits. The composite trait describing symbiont effects on resource acquisition arguably reflects the true currency of the host-symbiont interaction (Ankrah and Douglas 2018), its raw effect on potential fitness (Moran, Ochman and Hammer 2019). How this currency is used by the host is described by the resource allocation composite trait. As we have seen, allocation is not only controlled by the host but also seems to depend on symbiont features that yet remain to be elucidated. The simple bi-variate framework we developed here hence revealed the true dimensionality of symbiont effects. As for more complex multi-variates analyses of traits constrained by genetics or physiology (Blows *et al.* 2015; Collet and Fellous 2019), synthetizing multi-dimensional variation into fewer dimensions unveiled the true causality of phenotypic variation.

### Framework prospects and limits

The simple method we used borrows conceptual tools to multivariate quantitative genetics (Blows *et al.* 2015). It was made possible by established knowledge of physiological constrains leading to a trade-off between fitness components, namely speed of larval development and adult size (Stearns and Koella 1986; Nunney 1996; DeLong and Hanley 2013; Teder, Vellau and Tammaru 2014). Technically, the method shares principles with analytical frameworks developed in sexual selection studies (Berger *et al.* 2014). Like other quantitative genetics approaches, it requires the simultaneous phenotyping of numerous variants – here 18 symbiont strains in place of genotypes, inbred lines or crosses – which is not always possible. The question of whether symbionts affect host resource acquisition or allocation could be studied in many other fashions. Functional approaches that identify the molecular and physiological mechanisms of host-symbiont interaction are one obvious possibility. Experimental evolution can also reveal genetic (or symbiotic) constrains between traits and functions and evolved patterns can be compared to those of wild organisms (Fellous *et al.* 2014). Each of these solutions has its benefits and may be combine for completeness. The framework we propose, here, because it involves the comparison of a large number of symbiotic strains, allows drawing general conclusions regarding the effect of a class of symbionts, here yeast, on the focal host.

In the future, we will work towards improving the method in two main aspects. First, we will consider constrains between more than two fitness components. Even though most trade-offs documented in the literature involve pairs of traits, it is obvious that resources may be allocated to more than two functions. A forthcoming challenge will therefore be to increase the dimensionality of our method. It will necessitate to define the covariance between n traits under fixed conditions of resource availability. These multi-dimensional envelopes describing the n traits controlled by the same limiting resources will define resource allocation. Variation orthogonal to these envelopes may then reflect resource acquisition. The above line of reasoning points to the second challenge we will need to tackle: what are the actual shapes of the bi- or multi-dimensional trade-offs? Here, with Ockham’s razor logic we used a linear fit. In fact, we defined the trade-off shape as a linear fit orthogonal to estimated resource acquisition line. We had to follow this procedure because in our dataset variation in resource acquisition largely exceeded that of resource allocation. Ideally, trade-off shapes would be determined independently from the datasets to be analyzed. This reveal a challenging task as variation in resource acquisition is pervasive in empirical studies (Van Noordwijk and De Jong 1986; De Jong and Van Noordwijk 1992). One solution may be to use artificial selection on populations of similar genetic composition, so that all resource acquisition variation is shared, and selecting in different directions along trade-off envelope. An alternative may be to use clones in highly controlled setups and vary resource richness in order to investigate the direction and shape of resource acquisition variation in the considered phenotypic space. Note that hosts may alter their resource allocation in function of resource availability (Descamps *et al.* 2016), symbionts effects would add to such natural variation.

## Conclusions

We introduced a new framework based on the analysis of a well-established trade-off between fitness components in order to partition symbiont simultaneous effects on host resource acquisition (*i.e.*, nutrient provisioning) and allocation (*i.e.*, host plasticity). This framework unveiled symbiont effects invisible when single phenotypic traits were analyzed individually. In particular, we found that yeasts, essential nutritional symbionts of *Drosophila* larvae, had similar effects on the resource acquisition of several species of flies, but that effects on resource allocation varied among host species. These differences not only shed light on the ecology of *Drosophila* flies but also illustrate the evolutionary importance of distinguishing between conceptually different influences of symbiosis on host phenotype.

## Acknowledgements

We thank Salomé Bourg and Natacha Kremer for critical reading of an earlier version of this work and for their encouraging comments for the next version. We also thank Marie-Pierre Chapuis, Nicolas Rode, Delphine Sicard and Christophe Vorburger for their helpful comments. We thank Laure Benoit, Maxime Galan, Kenza Qitout and Antoine Rombaut for their help during the experiment or for the molecular identification of the microbial isolates. We thank Paul Becher for providing several yeast isolates.

## Author contributions

R.G. and S.F. designed the experiment. R.G. and A.X. ran the experiment. R.G. and S.F. analyzed the data and wrote the manuscript.

## Data availability

For the next version of this work, the dataset will be available in the open data repository Zenodo.

## Supplementary Material

### SM 1. Growth of exogenous microorganisms to our experiment

First, we investigated the potential factors that may have influenced the growth of exogenous microorganisms. We used a nominal logistic model with ‘*yeast treatment*’, ‘*fly population*’ (nested in ‘*fly group*’), ‘*fly group*’, their interactions and ‘*block*’ as fixed factors. Yeast treatment did influence the growth of exogenous microorganisms (χ^2^ = 43.17, df = 18, p = 0.0008) (Figure S1).

**Figure S1.**
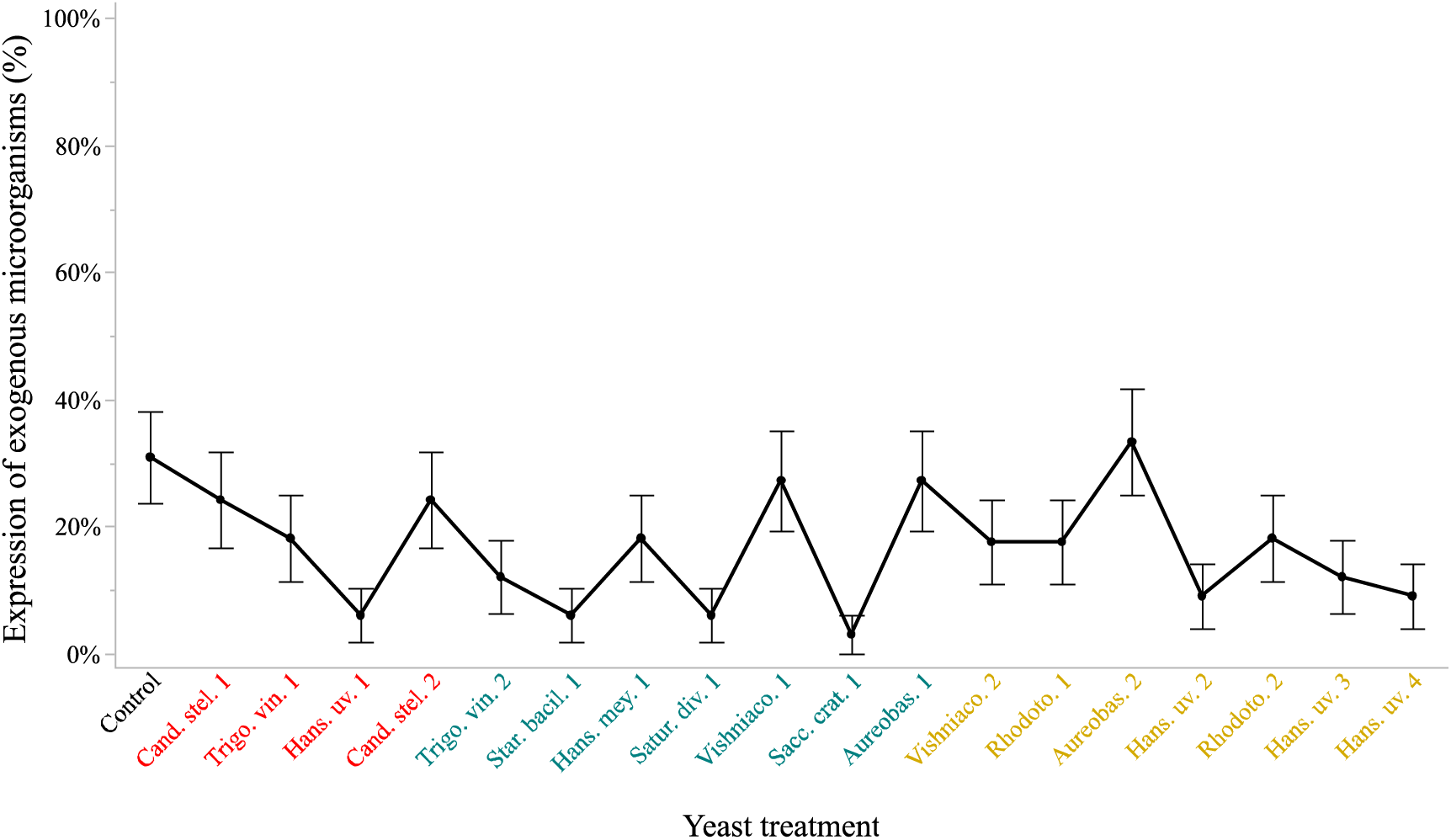
Growth of exogenous microorganisms per yeast treatment. Symbols indicate means and error bars indicate standard errors around the mean. X axis color code indicates the origin of the strains (red: *D. melanogaster*, blue: *D. suzukii*, green: fruit).

Second, we investigated whether these exogenous microorganisms may have affected fly individual traits (*i.e.*, developmental survival, age at emergence and thorax size). We used a linear mixed model (REML method) with ‘*absence or presence of exogenous microorganism*’, ‘*yeast treatment*’, ‘*fly population*’ (nested in ‘*fly group*’), ‘*fly group*’ and their interactions as fixed factors. ‘*Experimental block*’ (systematically) and ‘*experimental unit*’ (only for testing age and size) were defined as random factors. In general, the exogenous microorganisms did not affect significantly fly traits (but see marginally non-significant effects in Table S1). However, development survival was differently affected by these microorganisms among the fly populations (Table S1).

**Table S1.** Analysis of effects of exogenous microorganisms on individual traits. Linear mixed models (REML).

|  | **Developmental survival** | **Age at emergence** | **Thorax size** |
| --- | --- | --- | --- |
| Exogenous $\mu$ | $F_{1,584} = 3.64$; $p = 0.0570$ | $F_{1,565} = 3.05$; $p = 0.0813$ | $F_{1,115} = 0.73$; $p = 0.3944$ |
| Exogenous $\mu$ * yeast treatment | $F_{18,574} = 1.42$; $p = 0.1137$ | $F_{18,532} = 1.26$; $p = 0.2104$ | $F_{18,499} = 0.93$; $p = 0.5377$ |
| Exogenous $\mu$ * fly group | $F_{1,572} = 2.03$; $p = 0.1550$ | $F_{1,555} = 0.35$; $p = 0.5564$ | $F_{1,489} = 0.08$; $p = 0.7740$ |
| Exogenous $\mu$ * fly population | $F_{3,569} = 4.63$; $p = 0.0033$ | $F_{3,540} = 0.97$; $p = 0.4077$ | $F_{3,525} = 0.51$; $p = 0.6786$ |
| Exogenous $\mu$ * sex | | $F_{1,3254} = 0.29$; $p = 0.5932$ | $F_{1,2551} = 2.42$; $p = 0.1200$ |

### SM 2. Traits per population

**Figure S2.1.**
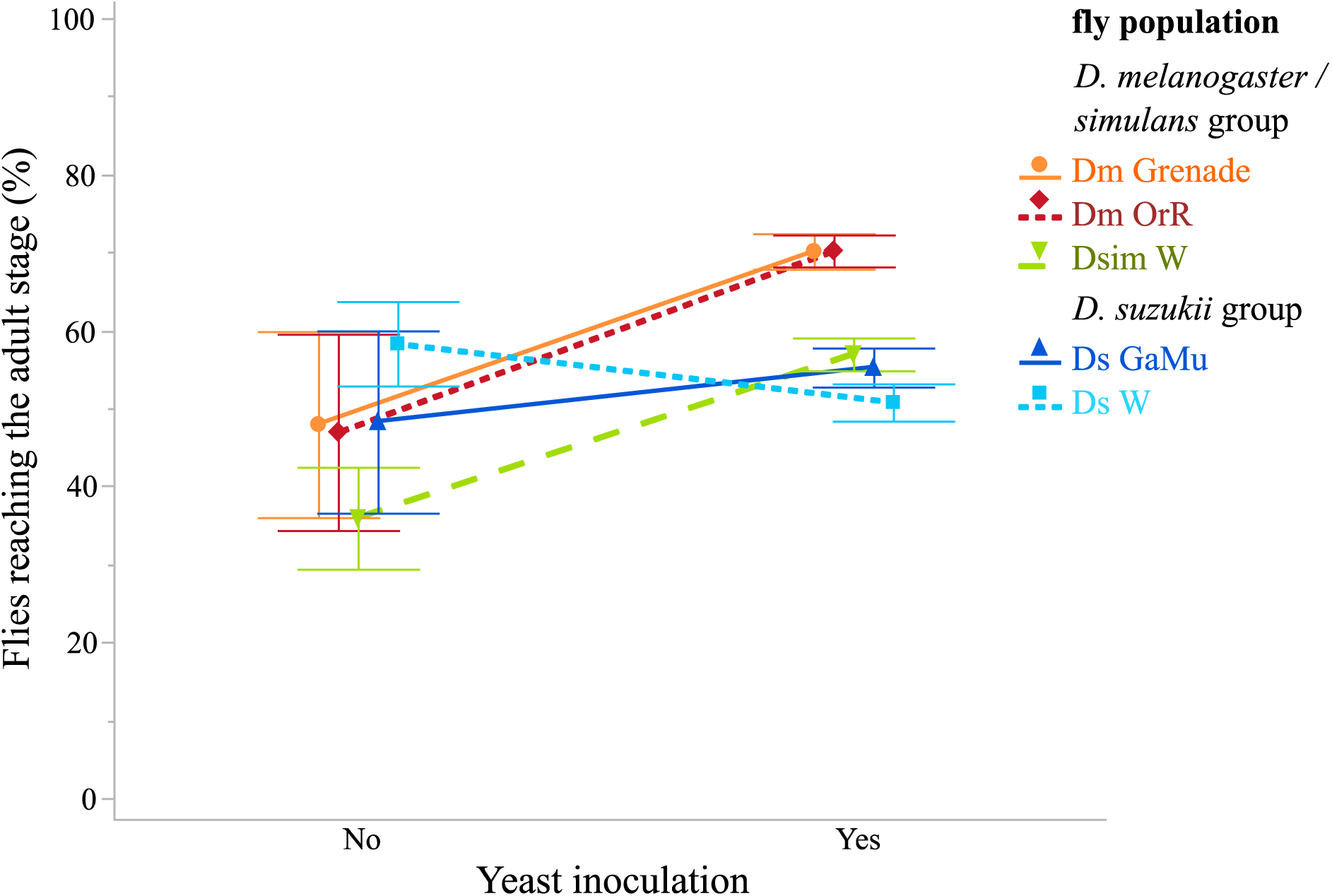
Development survival of each fly population in absence and presence of yeast. Symbols indicate means and error bars indicate standard errors around the mean.

**Figure S2.2.**
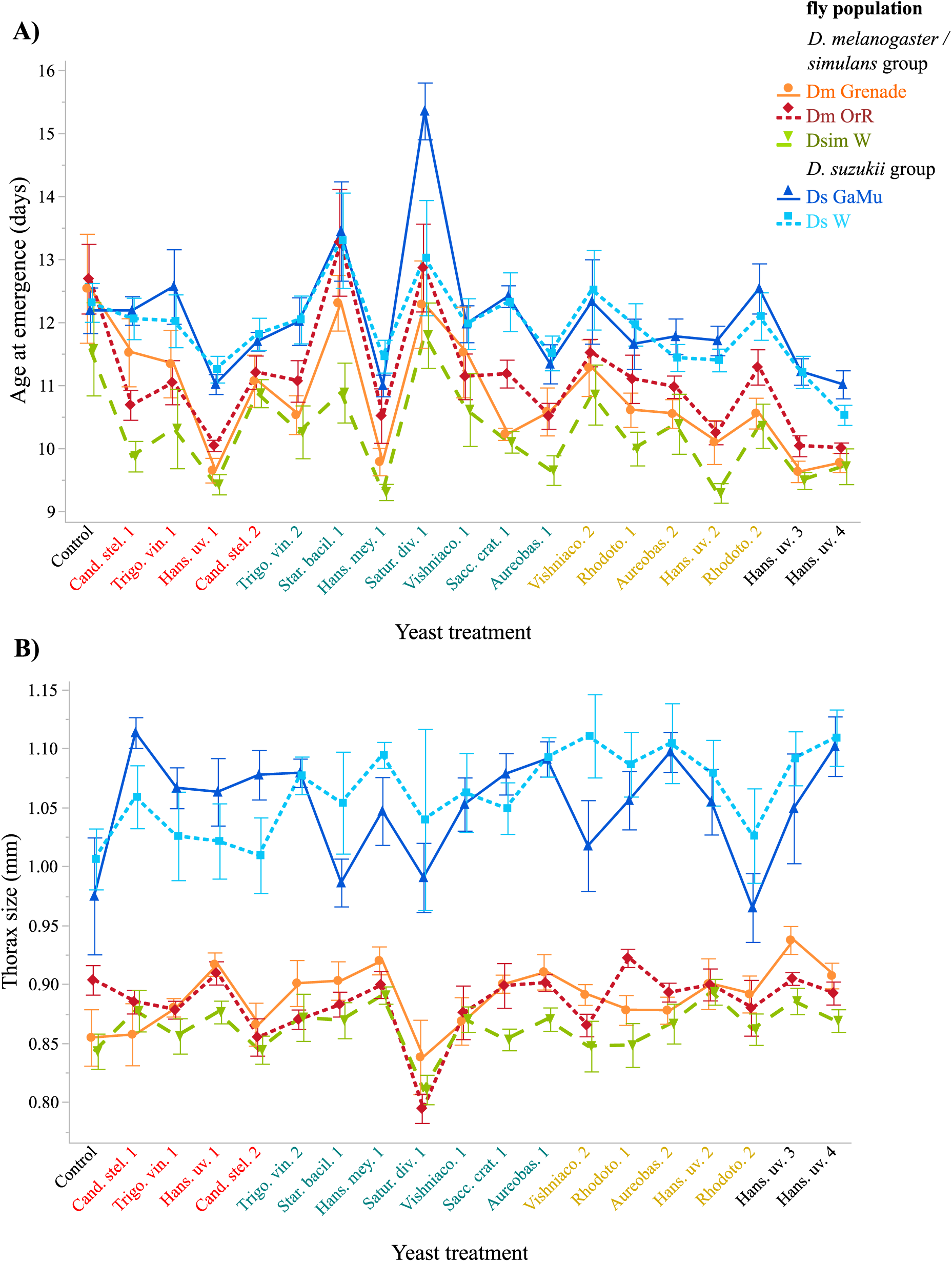
Age at emergence and thorax size of each fly population in response to yeast treatment. (A) Age at emergence per yeast treatment. (B) Thorax size per yeast treatment. Symbols indicate means and error bars indicate standard errors around the mean. X axis color code indicates the origin of the strains (red: *D. melanogaster*, blue: *D. suzukii*, green: fruit).

### SM 3. Relationships between yeast effects on speed of development and thorax size (symmetrical orthogonal regressions)

**Figure S3.**
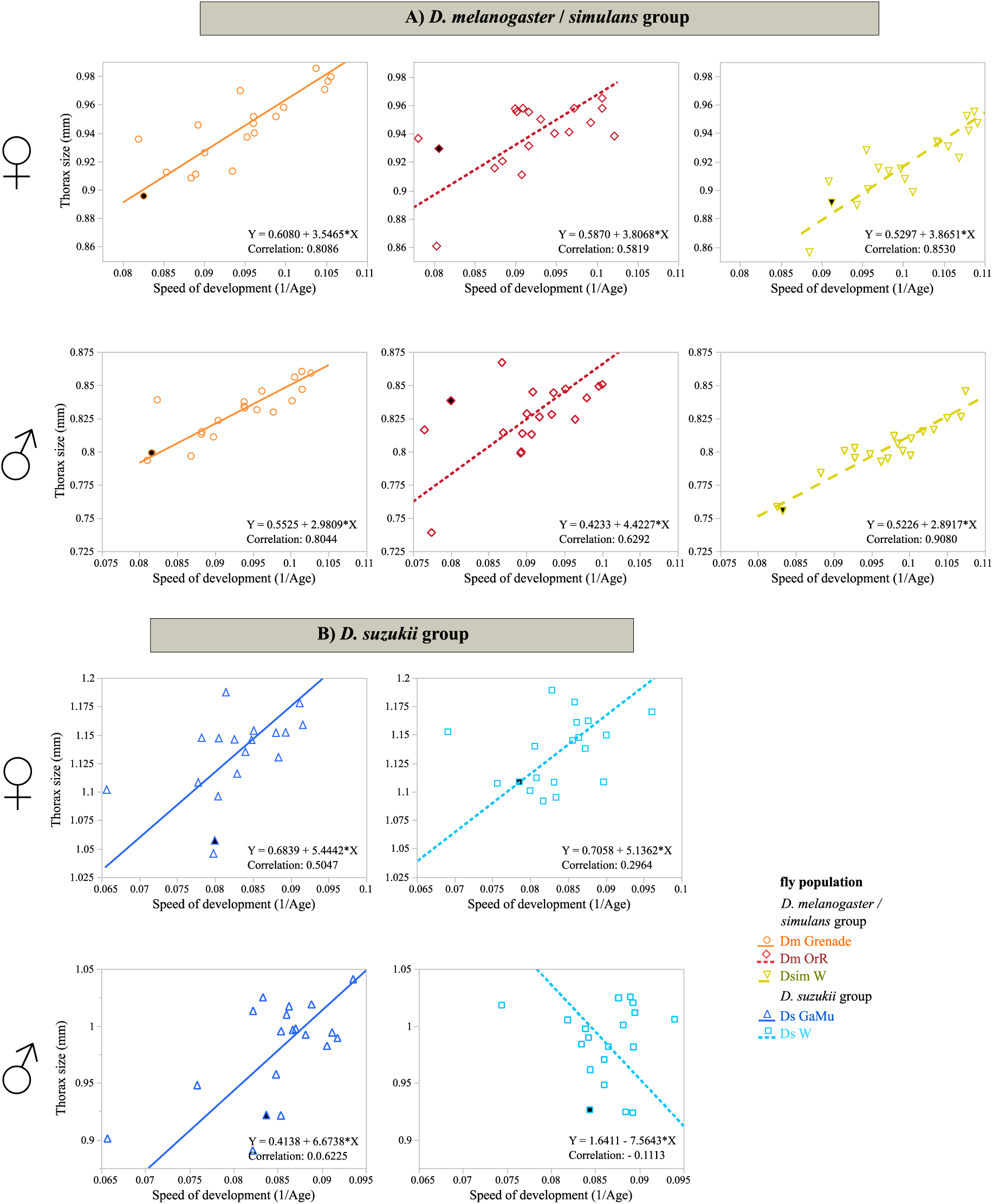
Relationships between yeast effects on larval speed of development and adult thorax size (symmetrical orthogonal regressions). Each point in the phenotypic space represents the mean of the two life-history traits affected by one yeast strain. Control phenotypes (*i.e.*, without yeast symbionts) are indicated by black-filled symbols.

### SM 4. Relationship between new response variables (resource acquisition and allocation) and original variables

**Figure S4.**
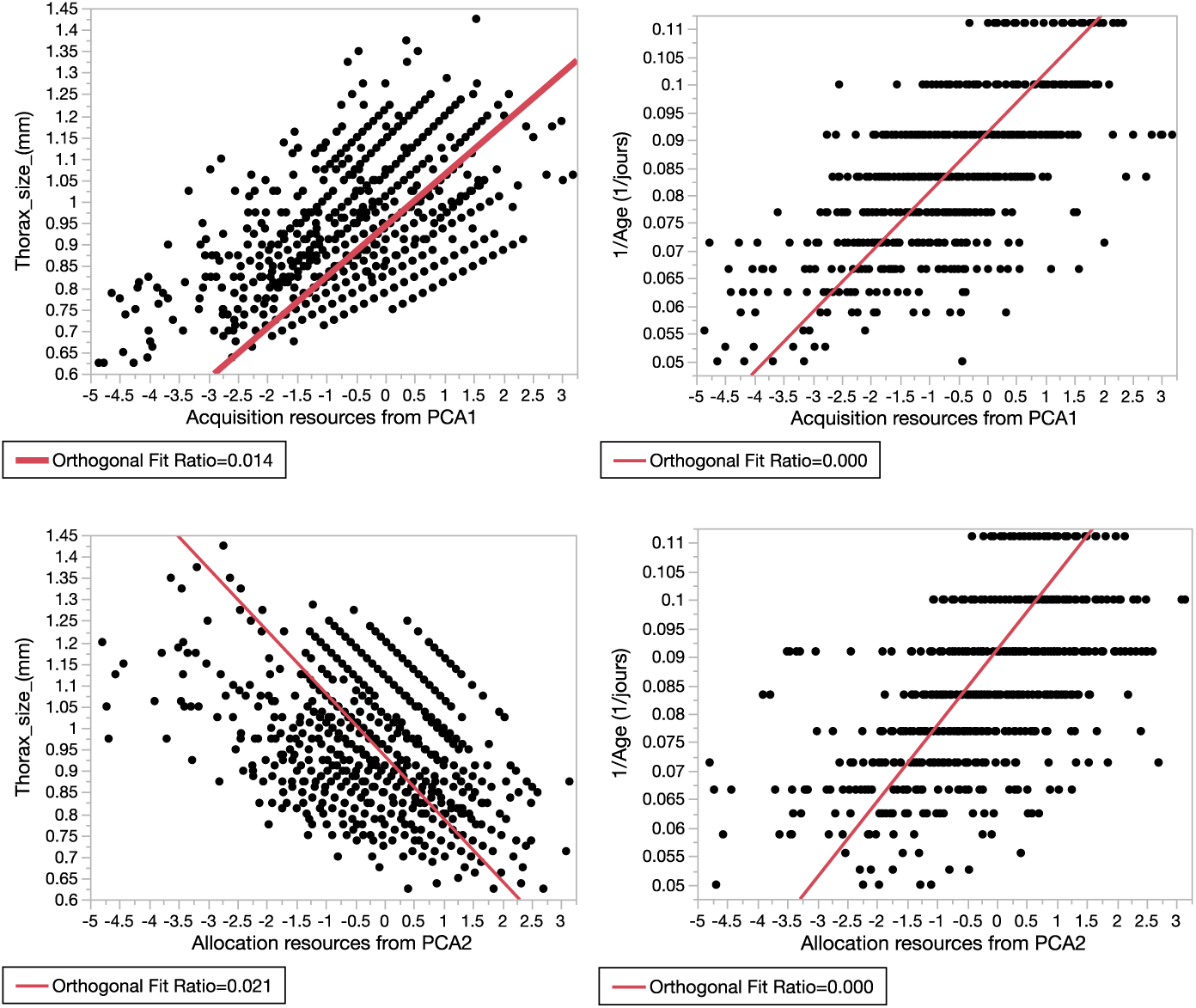
Relationship between new response variables (resource acquisition and allocation) and original variables.

### SM5. PC2 per population

**Figure S5.**
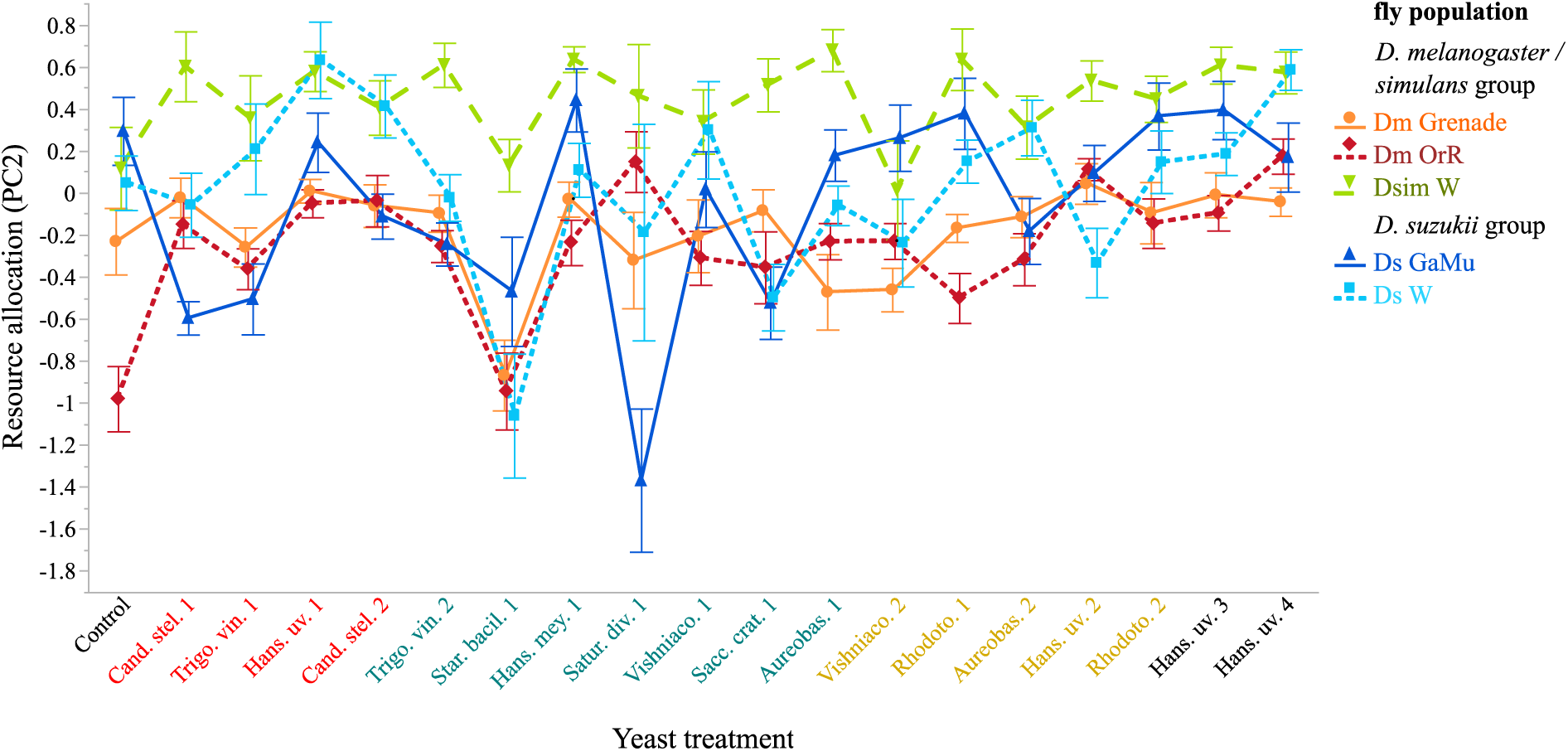
Resource allocation of each fly population in response to yeast treatment. Symbols indicate means and error bars indicate standard errors around the mean. X axis color code indicates the origin of the strains (red: *D. melanogaster*, blue: *D. suzukii*, green: fruit).

### SM6. Relationships between yeast density and phenotypic traits (OLS regressions)

**Figure S6.1.**
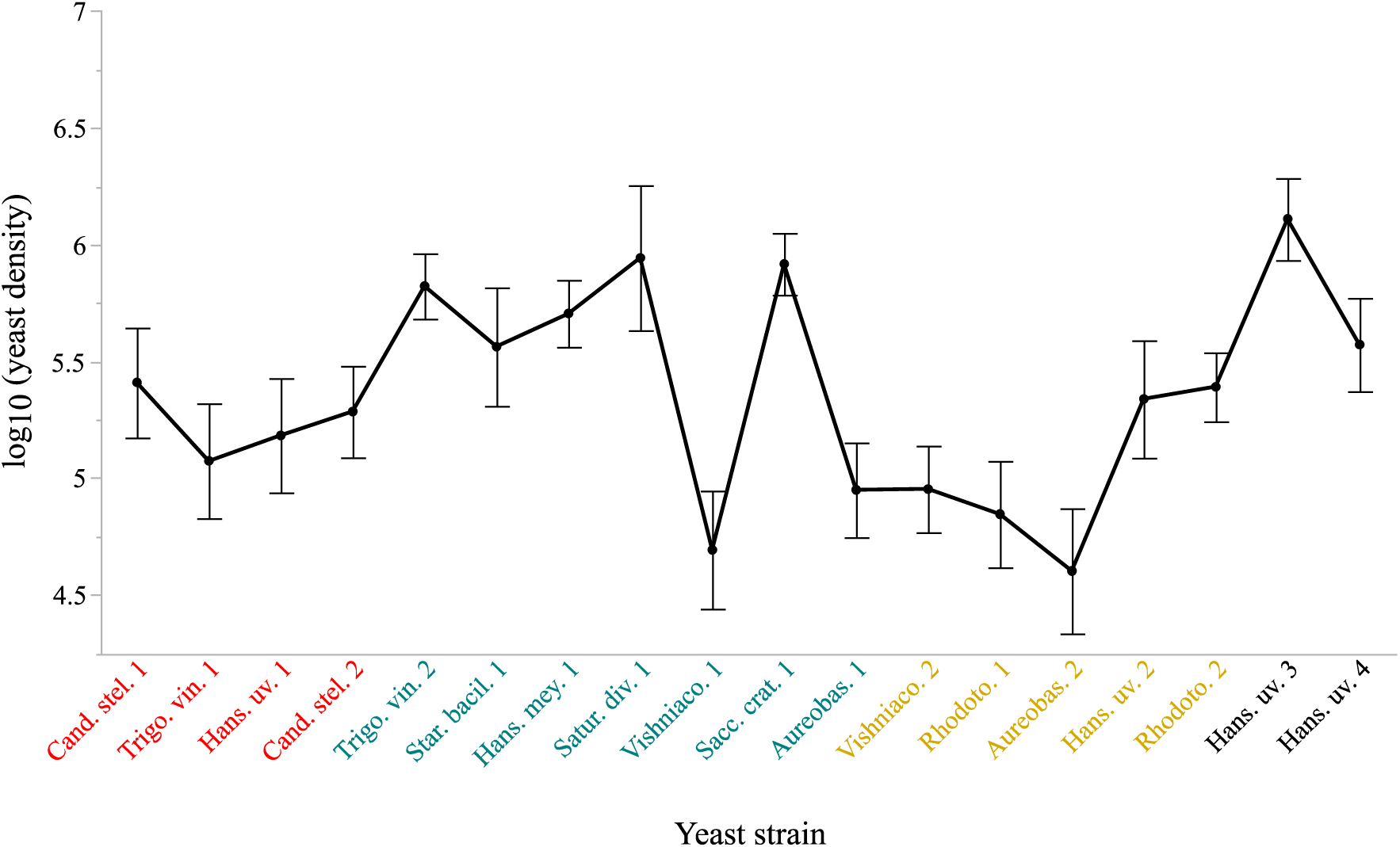
Yeast density in fruit flesh in response to yeast strain. Symbols indicate means and error bars indicate standard errors around the mean. X axis color code indicates the origin of the strains (red: *D. melanogaster*, blue: *D. suzukii*, green: fruit).

**Figure S6.2.**
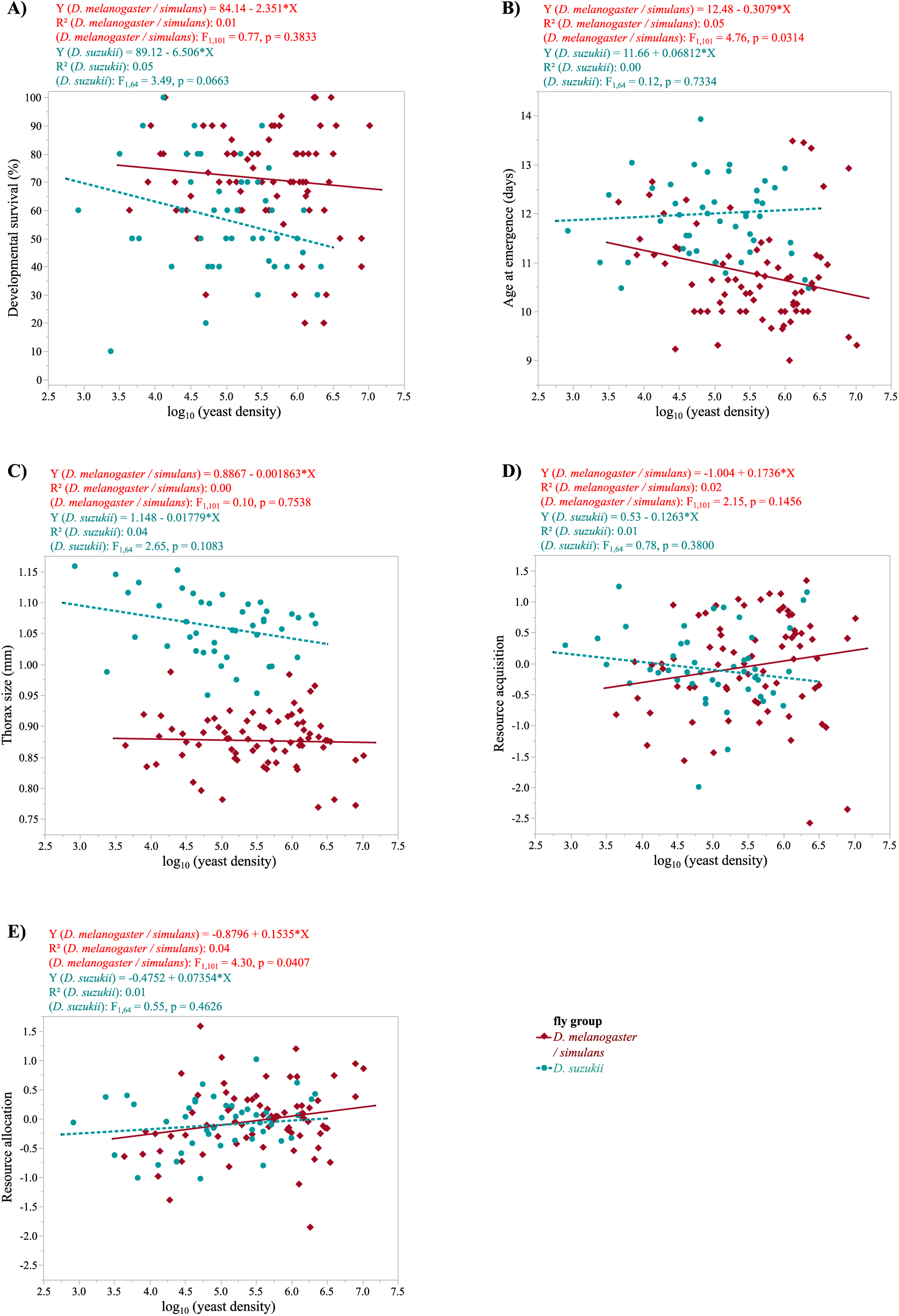
Linear regression between mean yeast density and mean fly traits. (A) Developmental survival. (B) Adult age at emergence. (C) Adult thorax size at emergence. (D) Resource acquisition. (E) Resource allocation.

### SM7. Effect of yeast origin on its influence of host development

**Table S7.**
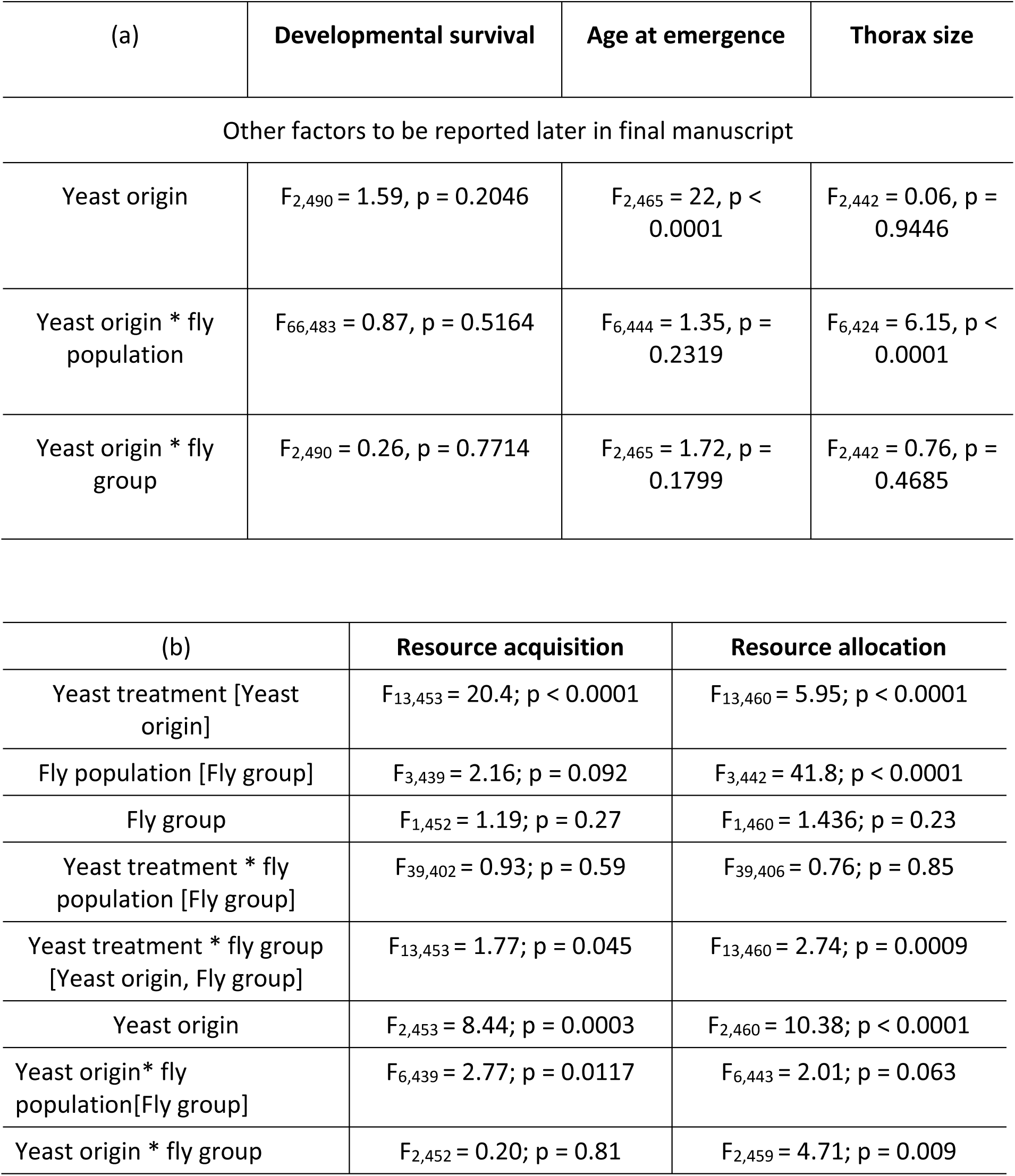
Analysis of effects of yeast origin on phenotypic traits. Linear mixed models (REML) of single traits (a) and new composite variables (b).

